# MIR17HG Expression Is Transcriptionally Regulated by PAX3::FOXO1 and MYCN and is Necessary for Oncogenic Activity in Fusion-Positive Rhabdomyosarcoma

**DOI:** 10.1101/2025.11.21.689335

**Authors:** Shabir Zargar, Pawan Kumar Raut, Hana Kim, Rachel A. Hoffman, Benjamin Z. Stanton, Frederic G. Barr

## Abstract

Alveolar rhabdomyosarcoma (RMS), an aggressive pediatric soft tissue cancer, is driven by the oncogenic fusion transcription factor PAX3::FOXO1 (P3F) or PAX7::FOXO1. In a subset of fusion-positive (FP)-RMS cases, amplification of the *MIR17HG* locus leads to overexpression of the miR-17-92 cluster of microRNAs (miRNAs). However, miR-17-92 is also highly expressed in FP-RMS tumors lacking this amplification, suggesting alternative regulatory mechanisms. Here, we show that P3F and MYCN cooperatively drive miR-17-92 expression in FP-RMS. CRISPR/Cas9-mediated knockout of *P3F* or *MYCN* in FP-RMS cell lines substantially reduced miR-17-92 expression. Using a human myoblast line or low P3F FP-RMS variant with inducible P3F or MYCN expression, P3F or MYCN alone induces minimal to low miR-17-92 expression whereas introduction of both MYCN and P3F leads to robust activation of the miR-17-92 cluster and acquisition of oncogenic phenotypes. Chromatin immunoprecipitation sequencing (ChIP-seq) revealed a P3F binding motif located 1.84 Mb upstream of the *MIR17HG* promoter. CRISPR-mediated deletion of this region in the myoblast system resulted in marked reduction of miR-17-92 expression and impaired oncogenic transformation. Functional inhibition of mature miRNAs of this cluster in FP-RMS cells using miRNA-sponge constructs suppressed proliferation and transformation. In the myoblast model system, transduction studies with exogenous miR-17-92 or miRNA-sponge expression constructs indicated that miR-17-92 is necessary but not sufficient for oncogenic transformation. Together, these findings establish a cooperative transcriptional axis in FP-RMS involving P3F and MYCN that activates *MIR17HG* through a distal regulatory element, thereby contributing to oncogenic behavior and uncovering a novel mechanistic vulnerability.

## Introduction

Rhabdomyosarcoma (RMS) is a pediatric soft tissue sarcoma, histologically classified into embryonal and alveolar subtypes. The alveolar subtype is characterized by recurrent chromosomal translocations, most commonly t(2;13) and t(1;13), which give rise to the *PAX3::FOXO1* (P3F) (1) and *PAX7::FOXO1* (P7F) fusion genes (2). These fusion genes encode novel transcriptional regulators that rewire the transcriptional machinery to promote proliferation and survival while blocking myogenic differentiation, ultimately driving the development of aggressive tumors (3,4). Due to the importance of these fusion oncoproteins, this RMS subtype is often referred to as fusion-positive RMS (FP-RMS).

MicroRNAs (miRNAs) are non-coding RNAs measuring approximately 22 nucleotides that regulate gene expression post-transcriptionally by promoting mRNA degradation or inhibiting translation (5). These small RNAs influence essential cellular processes, including proliferation (6), apoptosis (7) and differentiation (8), and are frequently dysregulated in cancer (5,9). Our laboratory has previously found that amplification of the 13q31 chromosomal region in FP-RMS encompasses the *MIR17HG* locus, whose primary transcript includes the oncogenic miR-17-92 cluster (miR-17, miR-18a, miR-19a, miR-20a, miR-19b-1 and miR-92a) (10). While this amplification is observed in a subset of FP-RMS, more commonly in P7F-positive tumors, elevated miR-17-92 expression is also detected in non-amplified FP-RMS tumors. These findings suggest that additional regulatory mechanisms contribute to *MIR17HG* activation beyond gene amplification (11). In neuroblastoma and medulloblastoma, *MYCN* gene amplification directly drives *MIR17HG* transcription (12). Since *MYCN* is a downstream target of P3F in FP-RMS (13), we hypothesize that P3F, either directly or through MYCN, drives miR-17-92 expression in FP-RMS.

This study investigates the upstream regulatory mechanisms and downstream phenotypic function of *MIR17HG* and the miR-17-92 cluster in FP-RMS. Using genetic perturbation (CRISPR-Cas9 knockout, inducible expression systems), functional assays and miRNA-sponge constructs, we explore the impact of P3F and MYCN and their interplay on expression and the oncogenic potential of miR-17-92 in FP-RMS cells. The findings reveal a critical oncogenic regulatory axis involving P3F, MYCN and MIR17HG, providing new insight into FP-RMS pathogenesis. This axis results in sustained miR-17-92 expression in FP-RMS, regardless of *MIR17HG* amplification status, and underscores the therapeutic relevance of targeting this pathway.

## Materials and Methods

### Cell Culture and Mycoplasma Testing

The Duchenne muscular dystrophy myoblast cell line (Dbt) and human FP-RMS cell lines were cultured under standard conditions. The sources of the cell lines are as follows: Dbt (D. Trono), Rh30 (ATCC), Rh30 Low P3F (E. Douglass), Rh28 (B. Emanuel), Rh5 (J. Khan) and Rh41 (C. Linardic). Rh30, Rh41, Rh28 and Rh30 Low P3F were maintained in Dulbecco’s Modified Eagle Medium (DMEM) supplemented with 10% fetal bovine serum (FBS). Rh5 cells were cultured in RPMI-1640 with 10% FBS, while Dbt cells were maintained in a 1:1 mixture of F-10 and DMEM supplemented with 15% FBS. All cultures were supplemented with penicillin-streptomycin (Gibco). Cell line identity was verified using short tandem repeat genotyping analysis with the AmpFLSTR Profiler Plus PCR Amplification Kit (Applied Biosystems). To rule out Mycoplasma contamination, cell lines were routinely tested using a PCR-based Mycoplasma detection kit (ATCC).

### Plasmid Constructs

To generate a constitutive *MYCN* expression construct, the *MYCN* open reading frame (ORF) was amplified from a previously cloned retroviral vector and inserted into the lentiviral backbone LeGO-iT (Addgene plasmid #27361). Directional primers were used for amplification, and the final construct was confirmed by Sanger sequencing. The miR-17-92 expression construct, containing the endogenous cluster of 6 miRNAs, was generated by amplifying the intronic region of *MIR17HG* from genomic DNA and cloning into a lentiviral expression plasmid (pLenti) containing a blasticidin resistance gene. For inducible expression of P3F and MYCN, we used the Dox-inducible lentiviral vector pInducer10b (Addgene plasmid #164935). The *P3F* ORF with a C-terminal 3×FLAG tag and the *MYCN* ORF with an N-terminal 3×FLAG tag were amplified from previously validated constructs. These inserts were cloned into the pInducer10b backbone using In-Fusion cloning (Takara Bio) and NEBuilder HiFi DNA Assembly (NEB). The resulting constructs, pInducer10b-P3F-3xFLAG-Puro and pInducer10b-3xFLAG-MYCN-BSD, were verified by sequencing.

### Design and construction of miRNA-Sponges

miRNA-sponges that functionally inhibit miRNAs of the miR-17-92 cluster were designed using the miRNAsong online tool (https://www.med.muni.cz/histology/miRNAsong/) (14). The binding sites in the sponge were reverse complements of the mature miRNA sequences with 3-4 mismatches outside the seed region. The sponge constructs were synthesized (GenScript) with six bulged binding sites for each targeted miRNA. Binding specificity was validated using the RNAhybrid tool (15). Two sponge sets targeting multiple miRNAs were constructed (16). SET1 targets miR-17, miR-20a and miR-92a and SET2 targets miR-18a, miR-19a and miR-19b1. SET1 and individual sponges for miR-17 and miR-92a were cloned into pInducer10b with a puromycin resistance gene. SET2 and the miR-20a sponge were cloned into the same backbone with the puromycin resistance gene replaced by a blasticidin resistance cassette. All constructs were verified by sequencing.

### Protein extraction, SDS-PAGE and Western Blotting

Cells were lysed in 1× SDS lysis buffer (Bio-Rad), followed by sonication using a cycle of 3 seconds ON / 3 seconds OFF on ice for a total of 8 pulses, or until the lysates appeared clear. Protein concentrations were determined using the Pierce BCA Protein Assay Kit (Thermo Scientific) according to the manufacturer’s instructions. It should be noted that β-mercaptoethanol was not added prior to protein estimation and was only included at the time that samples were prepared for gel loading. Unless otherwise stated, 30 µg of total protein per sample were loaded onto pre-cast TGX SDS-PAGE gels (Bio-Rad). These pre-stained gels allowed real-time visualization of protein migration during electrophoresis. Proteins were then transferred to PVDF membranes, which were subsequently blocked in 3% non-fat dry milk in Tris-buffered saline for 1 hour at room temperature. Membranes were incubated with antibodies against FOXO1 (1:1,000, Cat# 2880; Cell Signaling Technology), MYCN (B8.4.B) (1:1,000, Cat# sc-53993, Santa Cruz), GAPDH (1:2,000, Cat# sc-25778; Santa Cruz Biotechnology) and Vinculin (1:5,000, Cat# V9131, MilliporeSigma).

### Transfection and Production of Lentiviral Particles

For the generation of lentiviral particles, plasmid DNA transfections were carried out using linear polyethyleneimine (PEI) in HEK293T cells. A 3:1 PEI to DNA ratio was optimized for high-efficiency transfection in HEK293T. Lentivirus particles were produced in 10 cm^2^ dishes with the following components: 6.0 mg pPax2, 3.0 mg pMD2 and 10.0 mg expression plasmid along with 60 mg PEI. Lentiviral transductions were performed in the presence of 1mg/ml polybrene.

### RNA Extraction, cDNA Synthesis and Real-Time PCR

Total RNA was isolated using TRIzol™ Reagent (Invitrogen) according to the manufacturer’s instructions. For mRNA expression analysis, cDNA synthesis was performed using random hexamer primers and Superscript IV reverse transcriptase (Life Technologies). The *MIR17HG* transcript levels were quantified using exon-exon junction primers from the *MIR17HG* ORF whereas pri-miR-17-92 levels were measured using primers from *MIR17HG* intron 3; both of these transcripts were normalized to 18S rRNA. miRNA expression was analyzed via stem-loop reverse transcription [28] followed by quantitative PCR (qPCR) using miRNA-specific forward and stem-loop specific universal primers. qPCR analysis of miRNA was performed using SYBR Green chemistry, and data were quantified by 2^^-ΔΔCT^ methods [29]. miRNA expression was normalized to RNU6 snRNA. All qPCR reactions were performed with 3 technical replicates. Primer sequences are listed in Supplementary Table 1.

### Cell Growth Analysis

Cell proliferation assays were conducted using the Incucyte live-cell imaging system (Sartorius). Cells were seeded in appropriate densities in 24-well plates, and images were acquired every 6 hours over a period of 6 days (17). Confluence was calculated using Incucyte’s integrated software based on phase-contrast imaging. Growth curves were generated from confluence measurements to assess proliferation dynamics following various genetic perturbations. All experiments were performed with three wells for each condition and repeated in at least three independent experiments for validation. For all experiments involving inducible expression constructs, expression was induced in the presence of 500 ng/ml Dox (18).

### Clonogenic and Focus Formation Assays

The clonogenic assay was performed to assess colony growth by seeding 400 cells in 6 cm dishes in conditioned medium, which was prepared by growing cells for 48 hours, collecting and filtering the medium through a 0.22 µm filter, and mixing 1:1 with fresh medium. After 20 days, the medium was removed and then cells were fixed with methanol for 10 minutes and then stained with Giemsa for 10 minutes. Plates were imaged using Image Lab software (Bio-Rad). The focus formation assay of oncogenic transformation was performed by seeding 3.5 × 10⁵ NIH3T3 cells per plate along with 400 test cells. After 3 weeks, cells were fixed and stained with Giemsa, and the number and area of foci were quantified using ImageJ software (19). Both assays were conducted in triplicate with empty vector as control. For all experiments involving inducible expression constructs, expression was induced in the presence of 500 ng/ml Dox.

### miRNA Sensor Assay

To evaluate the efficacy of the miRNA-sponges (20), we used the miR-sensor assay (20) in pmirGLO vector (Addgene plasmid #49379). Three binding sites with perfect complementarity for miR-17a and miR-92a were cloned into the 3′ UTR of the luciferase gene. Sponge-expressing cells were transfected with the sensor constructs using the Fugene reagent. After 48 hours, luciferase expression was detected by western blot.

### Statistical Analysis

Experiments were performed in triplicate, and data are presented as means ± standard deviation (SD) of the means. Differences between test and control groups were analyzed using Prism software (GraphPad). An unpaired t-test was used to compare two samples, while one-way analysis of variance (ANOVA) with multiple comparisons was applied when three or more samples were involved in a single experiment. P-values were calculated and are shown in the figures as described in the figure legends.

### Data Availability Statement

All data supporting the findings of this study are provided within the article and its Supplementary Materials. Sequencing datasets generated in this study have been deposited in dbGaP under accession number phs004381.v1.p1. Additional data is available from the corresponding author upon reasonable request.

## Results

### P3F and MYCN Influence Expression of the miR-17-92 Cluster

Given that MYCN transcriptionally activates *MIR17HG* in neuroblastoma (21) and medulloblastoma (22), we examined the expression levels of the miR-17-92 cluster across a panel of FP-RMS lines with variable expression of P3F and MYCN. We compared miR-17-92 expression in these FP-RMS lines to an immortalized human myoblast line (Dbt) and a spontaneous non-transformed subclone of Rh30 cells with very low P3F levels (Rh30 Low P3F). The FP-RMS cell lines (Rh5, Rh28, Rh30 and Rh41) with moderate to high expression of both P3F and MYCN, as confirmed by western blot analysis (Figure 1A), showed high expression of the miR-17-92 cluster (Figure 1B). In contrast, the Dbt and Rh30 Low P3F lines, which exhibit low expression of both P3F and MYCN, demonstrated markedly reduced levels of miR-17-92. These findings support the premise that the upregulation of miR-17-92 may be related to the presence of either P3F, MYCN or both.

**Figure 1.**
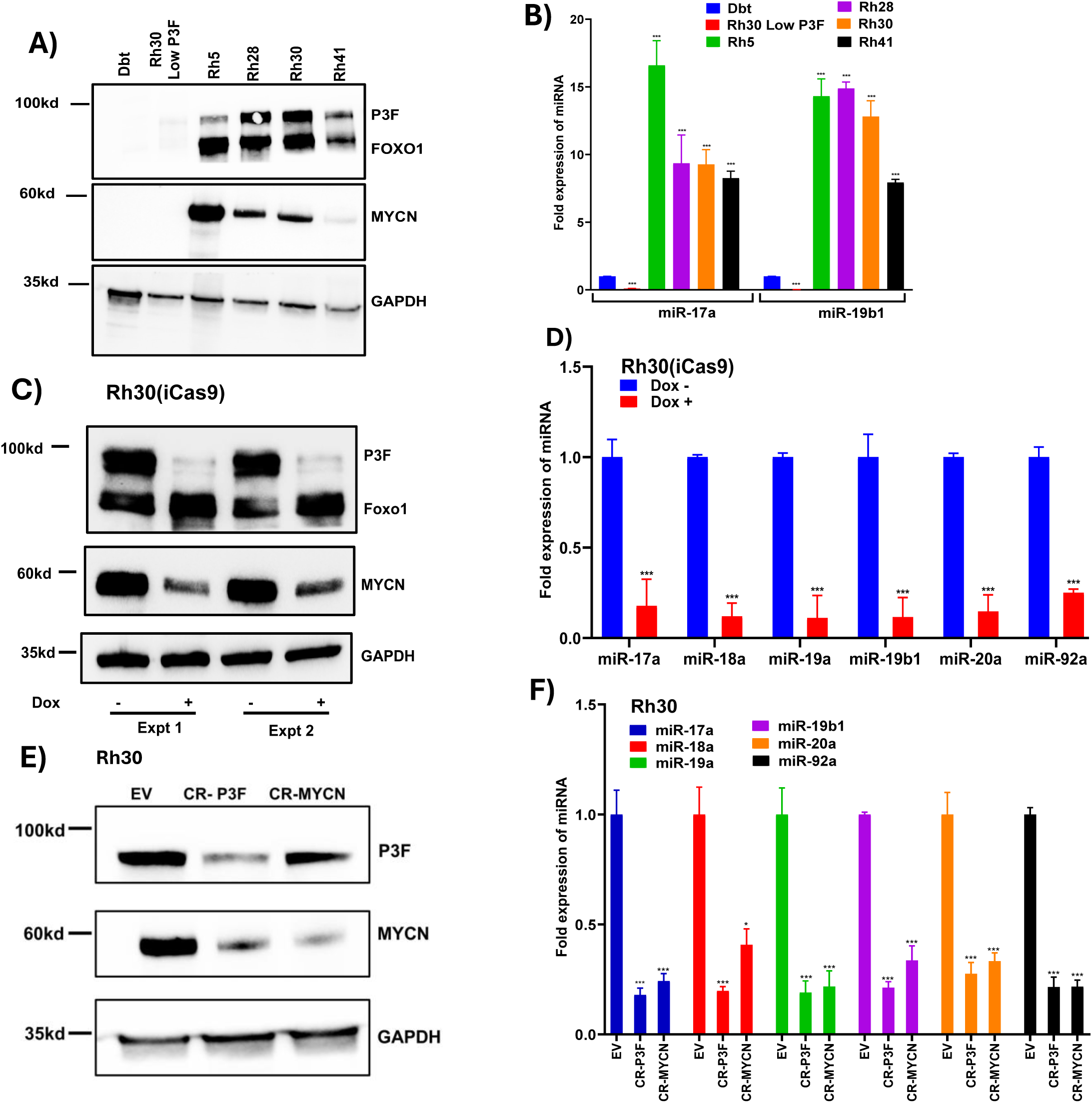
P3F and MYCN influence expression of the miR-17-92 cluster in FP-RMS. **A, B.** Western blot analysis of P3F and MYCN protein (A) and qPCR analysis of mature miR-17a and miR-19b1 expression (B) across a panel of FP-RMS cell lines (Rh5, Rh28, Rh30 and Rh41) and control lines (Rh30 Low P3F and Dbt). In A, GAPDH was utilized as a loading control. In B, results are normalized for expression in Dbt cells. **C, D.** Western blot of P3F and MYCN protein (C) and qPCR analysis of mature miRNAs within the miR-17-92 cluster (D) following doxycycline (Dox)-induced knockout of *P3F* in the Cas9-inducible Rh30 cells. Cells were treated without (-) or with (+) 2000 ng/ml Dox. In C, two independent experiments are shown. In D, results are normalized for cells without Dox. **E, F.** Western blot analysis of P3F and MYCN (E) and qPCR analysis of expression of mature miRNAs from the miR-17-92 cluster (F) following CRISPR-Cas9-mediated knockout (CR) of *P3F* or *MYCN* in Rh30 cells. Control cells were transduced with empty vector (EV). In F, results are normalized for these control cells. Data are presented as mean ± SD from at least three independent experiments. In B, D and F, statistical significance is displayed as: not significant (ns), p < 0.05 (*), p < 0.01 (**), p < 0.001 (***).

To investigate the role of P3F in regulating miR-17-92 expression, we utilized previously established Rh30 and Rh41 subclones that harbor a doxycycline (Dox)-inducible Cas9 system targeting P3F. CRISPR-Cas9-mediated knockout (KO) of P3F was induced by Dox treatment, and efficient loss of P3F protein was confirmed by western blot analysis in Rh30 (Figure 1C) and Rh41 (Figure S1A). P3F depletion resulted in a marked reduction in MYCN protein levels, consistent with *MYCN* being a downstream transcriptional target of P3F (13). qPCR analysis demonstrated that P3F KO led to a significant reduction in *MIR17HG* transcript and pri-miR-17-92 levels in both Rh30 (Figure S1B) and Rh41 (Figure S1C) cells. Correspondingly, the expression of mature miR-17-92 miRNAs was also markedly decreased in Rh30 (Figure 1D) and Rh41 (Figure S1D).

Given that MYCN is a known regulator of *MIR17HG* in neuroblastoma and medulloblastoma, we sought to determine whether MYCN is necessary for the miR-17-92 expression in these FP-RMS cells. To dissect the contribution of MYCN, we performed targeted MYCN knockout as well as targeted P3F KO using CRISPR-Cas9 in Rh30 and Rh41 cells. Western blotting confirmed efficient depletion of each protein in Rh30 (Figure 1E) and Rh41 (Figure S2A) and confirmed that MYCN expression is dependent on P3F in these FP-RMS cells with P3F KO. To systematically evaluate the impact on the miR-17-92 locus, we measured expression of MIR17HG transcript, pri-miR-17-92, and mature miRNAs. qPCR showed that loss of either P3F or MYCN significantly reduced mature miR-17-92 miRNA expression in Rh30 (Figure 1F) and Rh41 (Figure S2B), consistent with results obtained from the inducible Cas9 KO. Correspondingly, *MIR17HG* and pri-miR-17-92 transcript levels were also markedly decreased following P3F or MYCN depletion in both Rh30 and Rh41 (Figure S2C, S2D). These results indicate that MYCN is necessary for high-level miR-17-92 expression, and that the effect of P3F at least partially depends on its ability to upregulate MYCN.

### P3F and MYCN Cooperatively Induce miR-17-92 Expression

To define the individual and cooperative contributions of P3F and MYCN to the regulation of miR-17-92 expression, we introduced inducible expression constructs for P3F and/or MYCN into two independent models: immortalized Dbt human myoblasts and Rh30 Low P3F cells. In our human myoblast system with inducible P3F expression (Dbt-iP3F) (23), P3F expression is controlled by a Dox-inducible promoter and is not sufficient to upregulate high-level MYCN expression in these cells (Figure 2A). We transduced Dbt control cells or Dbt-iP3F cells with a lentiviral construct that constitutively expresses MYCN to generate cells expressing MYCN alone or P3F and MYCN. qPCR analysis revealed that P3F alone or MYCN alone elicited only a modest increase in miR-17-92 expression (Figure 2B). In contrast, co-expression of MYCN and P3F led to a marked elevation of miR-17-92 levels. These results suggest a synergistic interaction between MYCN and P3F in activating miR-17-92 expression.

**Figure 2.**
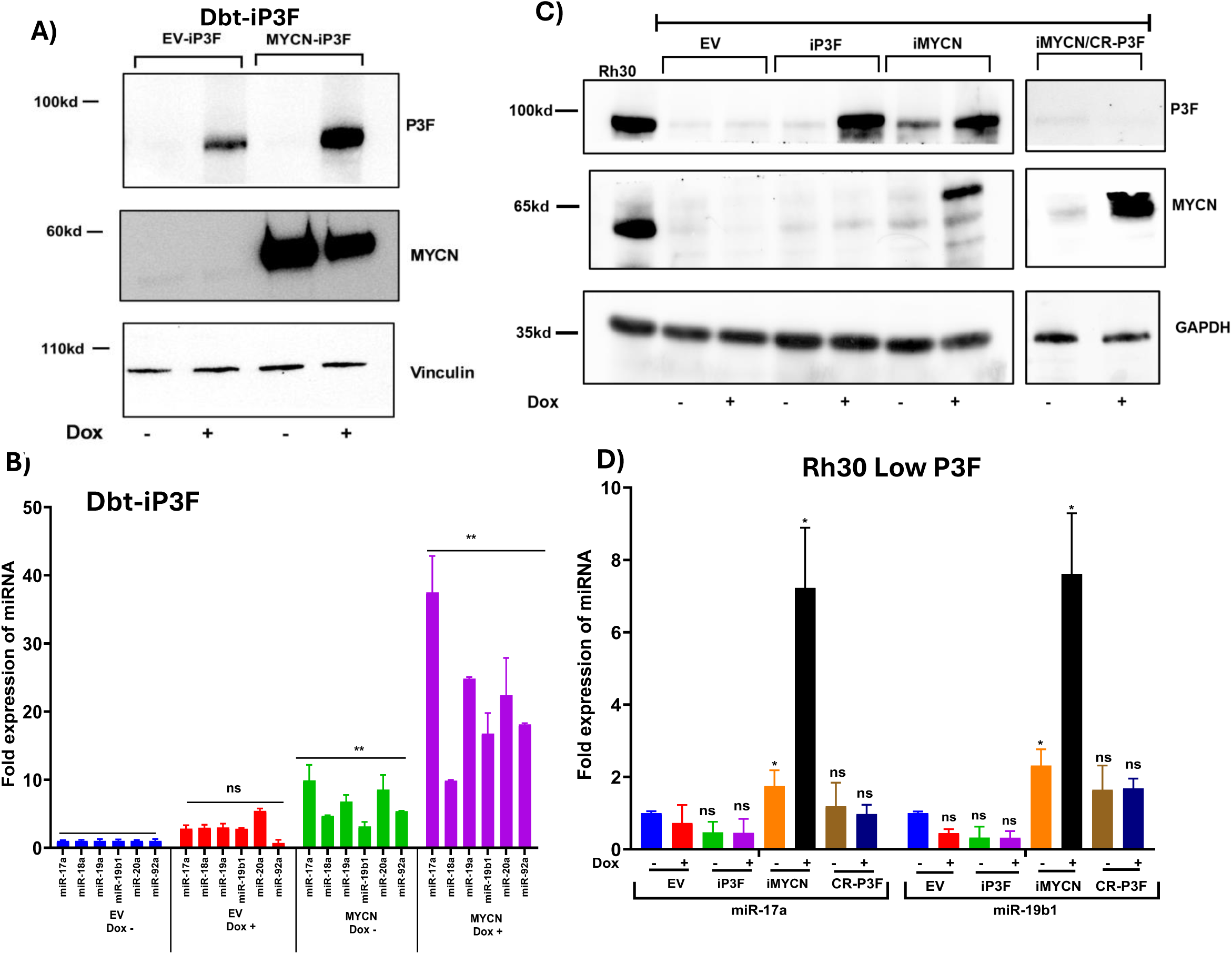
P3F and MYCN cooperatively induce miR-17-92 expression. **A, B.** Western blot analysis of P3F and MYCN protein (A) and qPCR analysis of miR-17-92 cluster expression (B) in Dbt myoblasts transduced with inducible P3F (iP3F) and/or constitutive *MYCN* construct and treated without (-) or with (+) 500 ng/ml doxycycline (Dox). In A, vinculin serves as a loading control. **C, D.** Western blot of P3F and MYCN protein (C) and qPCR analysis of miR-17a and miR-19-1 expression (D) in Rh30 Low P3F cells harboring iP3F or iMYCN inducible construct, or iMYCN with knockout of P3F (CR-P3F). Cells were treated without (-) or with (+) 500 ng/ml Dox. In C, GAPDH serves as a loading control. In B and D, results were normalized for EV-transduced cells without Dox. Data are presented as mean ± SD from at least three independent experiments. In B and D, statistical significance is displayed as described in Figure 1.

To validate this finding in a second model system related to FP-RMS, we used the Rh30 Low P3F cells to establish two Dox-inducible lines, one for MYCN (iMYCN) and another for P3F (iP3F). Upon Dox treatment, western blot confirmed robust induction of each transgene (Figure 2C). Interestingly, MYCN induction alone led to an increase in endogenous P3F expression, reaching levels similar to those observed in Rh30 FP RMS cells. In qPCR studies of miR-17-92 expression, induction of P3F alone did not upregulate miR-17-92 expression whereas MYCN induction (which also induces P3F expression) upregulated expression of the miR-17-92 cluster (Figure 2D). To dissect the requirement for P3F in MYCN-driven miR-17-92 expression, we employed a CRISPR/Cas9 approach to knockout P3F in the iMYCN cells (Figure 2C). In contrast to the significant miR-17-92 activation in the iMYCN cells (in which both MYCN and P3F are induced), MYCN induction in the P3F KO iMYCN cells failed to upregulate miR-17-92 (Figure 2D). These gain-of-function studies support a model in which P3F and MYCN act cooperatively and are sufficient to drive high level miR-17-92 expression in FP-RMS.

### Inhibition of the miR-17-92 Cluster Suppresses Oncogenic Properties in FP-RMS

To investigate the functional importance of the miR-17-92 cluster in FP-RMS, we employed a miRNA-sponge-based approach (24) to inhibit the activity of mature miRNAs derived from the MIR17HG locus. These synthetic sponge constructs contain tandem binding sites with partial complementarity to specific miRNAs, enabling them to sequester endogenous miRNAs and relieve repression of target mRNAs (25) (Figure S3A). For these experiments, we divided the six miRNAs of the miR-17-92 cluster into two sets, each designed to target three miRNAs. SET1 targets miR-17a, miR-20a and miR-92a whereas SET2 targets miR-18a, miR-19a and miR-19b-1. In addition, we generated individual sponges for miR-17a, miR-20a and miR-92a to further dissect the contributions of these miRNAs (Figure S3B). Of note, all these sponge constructs were cloned into Dox-inducible expression vectors.

We validated the functional efficacy of two miRNA-sponge constructs using a luciferase-based miRNA sensor assay (20). Sensor constructs containing three tandem binding sites with perfect complementarity to either miR-17a or miR-92a were cloned into the 3′ UTR of a luciferase reporter gene (Figure S3C). These constructs were transfected into Rh30 cells stably expressing the corresponding miRNA-sponge or control (empty vector). In empty vector (EV) controls, endogenous miRNAs bind to the sensors and suppress luciferase expression. In contrast, the miRNA-sponges sequester the corresponding miRNAs and relieves this suppression, resulting in increased luciferase protein detected by western blot analysis (Figure S3D). These findings verify that the sponges effectively inhibit miRNA activity and support their utility for downstream functional studies.

To assess whether miR-17-92 activity is essential for oncogenic transformation, we employed the Dbt-MYCN-iP3F model, in which MYCN is constitutively expressed and P3F is Dox-inducible, and we transduced Dox-inducible miRNA-sponge constructs into these cells. Upon Dox induction of P3F, EV-transduced cells formed dense colonies and foci, indicative of oncogenic transformation (Figure 3A, 3B) In contrast, cells with Dox induction of both P3F and either SET1 or individual sponges targeting miR-17a, miR-20a, or miR-92a showed markedly reduced transformation, as evidenced by fewer and smaller colonies and foci. In the clonogenic assay of cells transduced with miRNA-sponge constructs, the finding of decreased colony formation without Dox induction is attributed to leaky expression of the miRNA-sponges. Notably, cells transduced with the SET2 sponge construct, which targets miR-18a, miR-19a and miR-19ba, formed dense colonies and foci upon Dox induction, similar to EV controls (Figure 3C, 3D), indicating that miR-18a, miR-19a, and miR-19b-1 have little to no role in suppressing oncogenic transformation under these conditions. Live-cell imaging further demonstrated that Dox induction of sponges targeting SET1, miR-17a, and miR-20a significantly impaired proliferation, while the miR-92a sponge had a more modest effect (Figure 3E). These results indicate that miR-17a, miR-20a, and miR-92a are the principal effectors of this miRNA cluster that drive oncogenic phenotypes in this myoblast model system.

**Figure 3.**
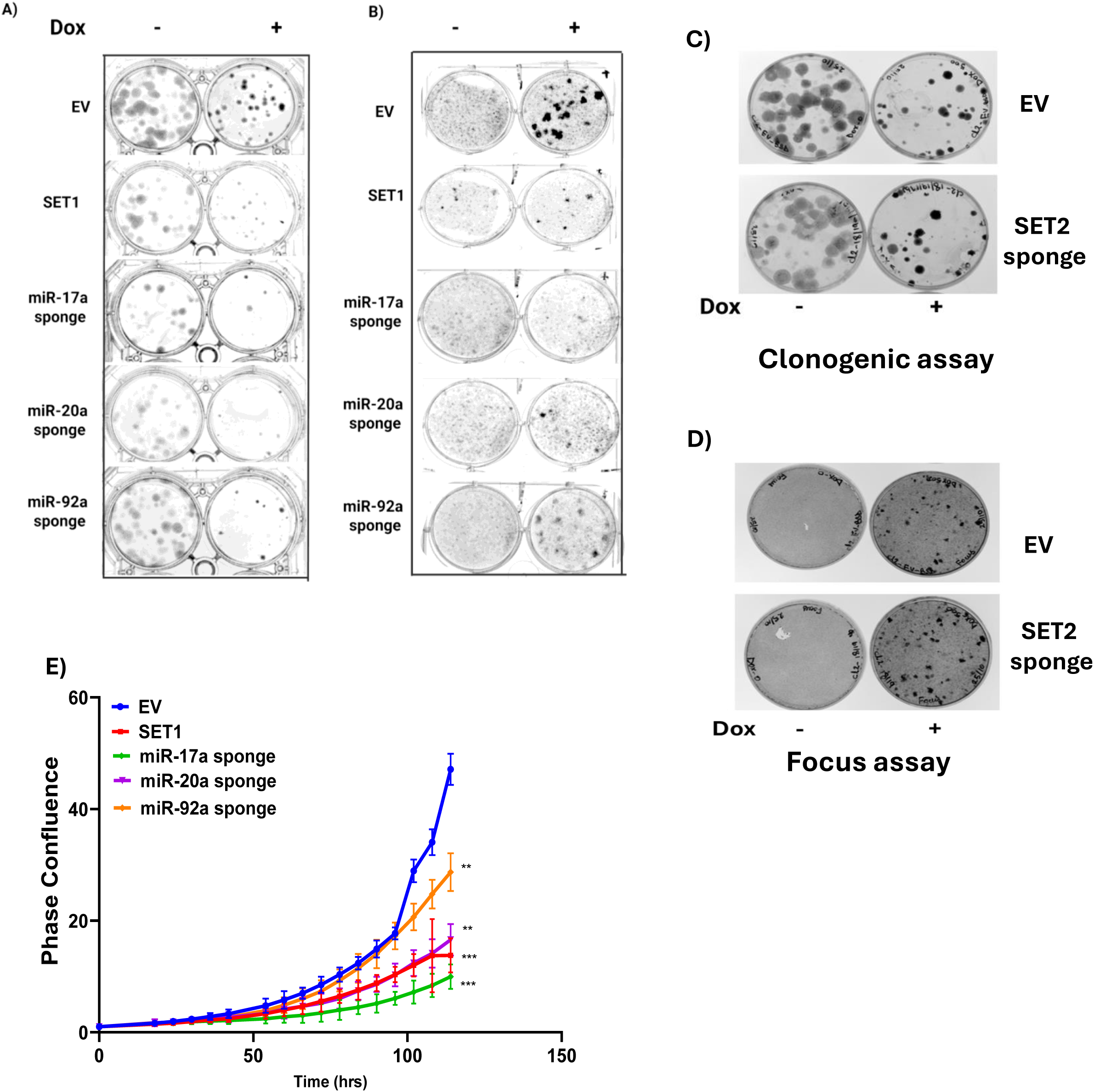
Inhibition of miR-17-92 impairs proliferation and transformation in myoblasts. **A, B.** Clonogenic (A) and focus formation (B) assays in Dbt-MYCN-iP3F cells expressing empty vector (EV), SET1 sponge or individual sponges for miR-17a, miR-20a and miR-92a and treated with (-) or without (+) doxycycline (Dox). **C, D.** Clonogenic (C) or focus formation (D) assays in Dbt-MYCN-iP3F cells expressing EV or SET2 sponge and treated with (-) or without (+) Dox. **E.** Proliferation analysis of Dbt-MYCN-iP3F cells expressing EV, SET1 sponge or individual sponges for miR-17a, miR-20a and miR-92a and treated with Dox. In E, statistical significance is displayed as described in Figure 1.

To determine whether this requirement for miR-17-92 extends to FP-RMS cell lines, we introduced SET1, SET2, individual miRNA-sponges or empty vector into Rh30 and Rh41 cells. Live-cell imaging of Dox-treated cells revealed distinct responses: in Rh30 cells, SET1 and SET2 sponges as well as the three individual sponges (from SET1) similarly suppressed proliferation, indicating a broad dependency on the miR-17-92 cluster (Figure 4A). In Rh41 cells, both SET1 and SET2 decreased proliferation, and for the three individual sponges from SET1, proliferation was selectively reduced by inhibition of miR-20a and miR-92a, but not miR-17a (Figure 4D). These findings point to the presence of variable cell line-specific dependencies on individual miRNAs within the cluster.

**Figure 4.**
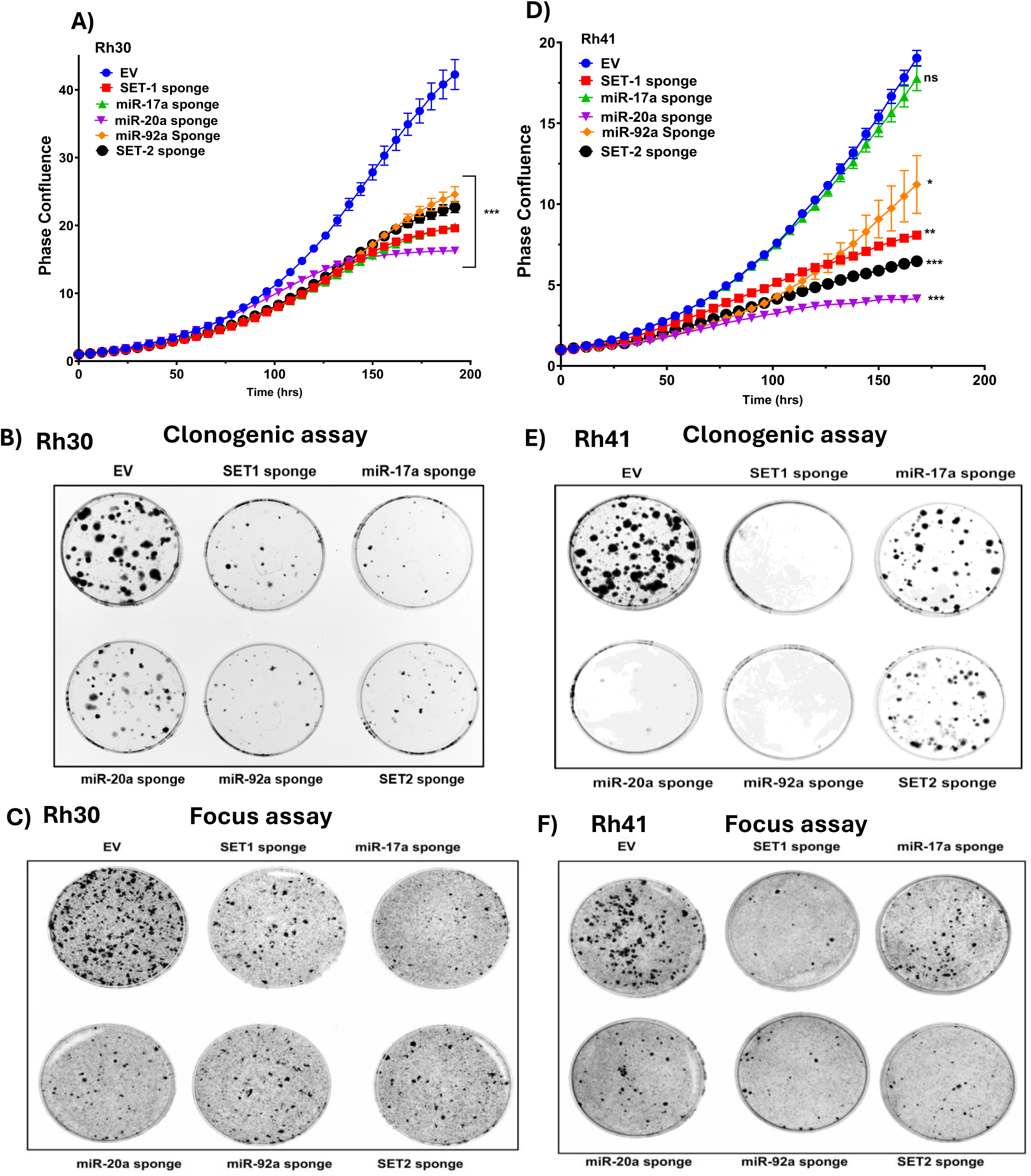
Inhibition of miR-17-92 suppresses proliferation and transformation properties in FP-RMS cells. **A.** Proliferation analysis of Rh30 cells expressing empty vector (EV), SET1, SET2, or individual sponges for miR-17a, miR-20a, and miR-92a. **B, C**. Clonogenic (B) and focus formation (C) assays in Rh30 cells expressing EV, SET1, SET2, or individual sponges for miR-17a, miR-20a, and miR-92a. **D.** Proliferation analysis of Rh41 cells expressing EV, SET1, SET2, or individual sponges for miR-17a, miR-20a, and miR-92a. **E, F**. Clonogenic (E) and focus formation (F) assays in Rh41 cells expressing EV, SET1, SET2, or individual sponges for miR-17a, miR-20a, and miR-92a. In A-F, cells were treated with 500 ng/ml doxycycline. Data are presented as mean ± SD from at least three independent experiments. In A and D, statistical significance is displayed as described in Figure 1.

Clonogenic and focus formation assays further supported these observations. In Rh30 cells, expression of SET1, its individual components or SET2 reduced colony and focus formation (Figure 4B, 4C). In Rh41 cells, similar reductions in colony and focus formation were observed with SET1 and sponges for miR-20a and miR-92a, while the sponges for miR-17a and SET2 only modestly impaired clonogenic growth, again indicating context-dependent roles for specific miRNAs (Figure 4E, 4F). The distinct clonogenic responses underscore the differential dependence of Rh30 and Rh41 cells on the component miRNAs of the miR-17-92 cluster.

### miR-17-92 Cluster is not Sufficient for Oncogenic Transformation

Using our human myoblast system, we next examined whether the miR-17-92 cluster alone can drive oncogenic transformation. A lentiviral expression construct was used to express the miR-17-92 cluster in Dbt myoblasts. Despite high expression of the pri-miR-17-92 compared to Rh30 FP-RMS cells (Figure S4A), these miRNAs did not enhance proliferation (Figure 5A); in fact, a modest reduction in cell growth was observed compared to empty vector-transduced control cells. Moreover, the Dbt myoblasts expressing the miR-17-92 cluster did not show an increase in the number or size of the colonies in a clonogenic assay (Figure 5B) and failed to exhibit focus formation, (Figure 5C). These findings indicate that miR-17-92 expression alone is insufficient to initiate oncogenic transformation.

**Figure 5.**
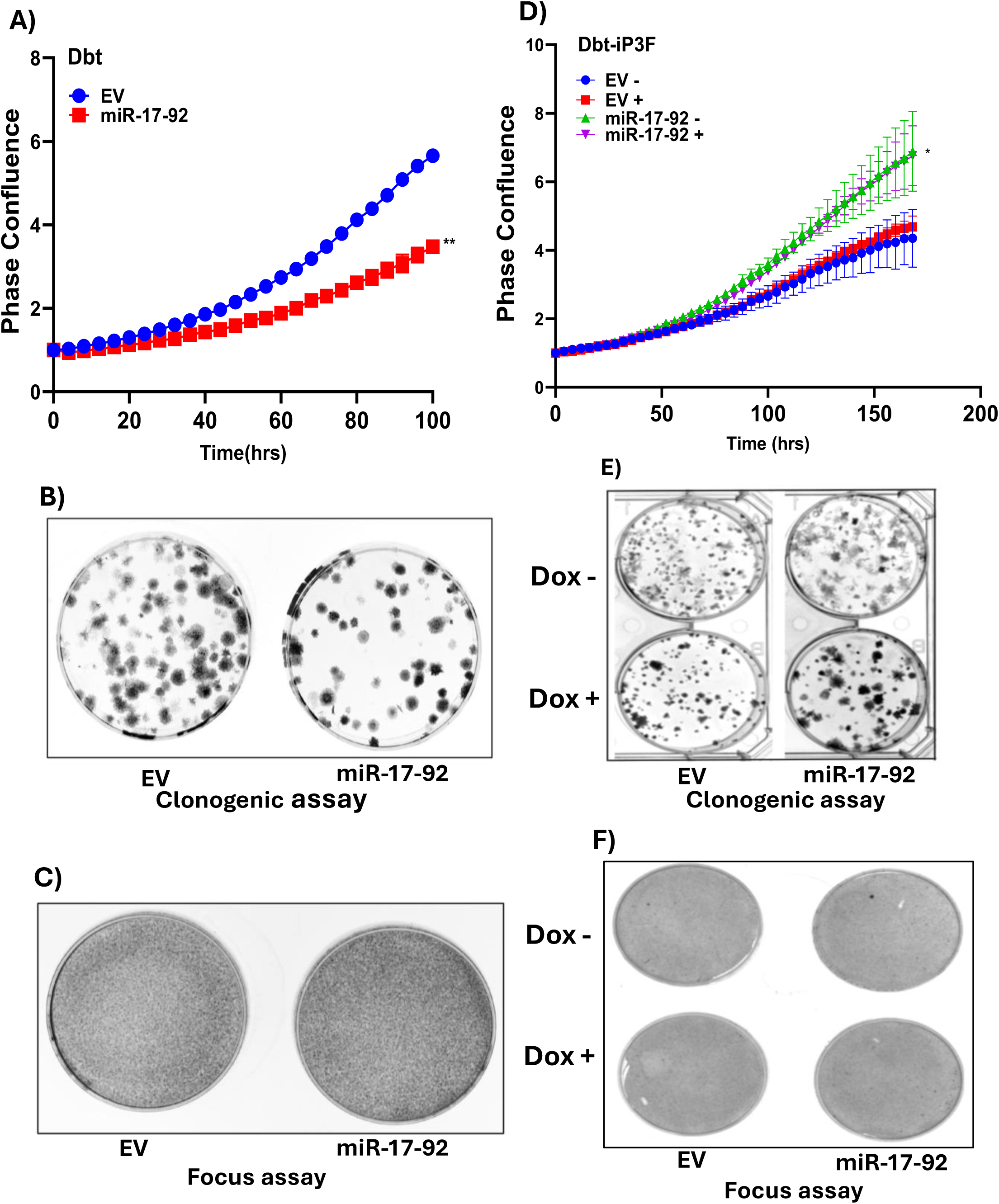
Phenotypic effects of miR-17-92 expression in human myoblasts. **A, B, C.** Proliferation (A), clonogenic (B) and focus formation (C) assays of Dbt myoblasts transduced with a lentiviral vector expressing the miR-17-92 cluster compared to empty vector (EV) control. **D, E, F.** Proliferation (D), clonogenic (E) and focus formation (F) assays of Dbt myoblasts with doxycycline (Dox)-inducible P3F transduced with lentiviral construct expressing miR-17-92 or EV control. In D, E, and F, cells were treated without (-) or with (+) 500 ng/ml Dox. In A and D, statistical significance is displayed as described in Figure 1.

Next, we assessed whether miR-17-92 could cooperate with the P3F oncoprotein in Dbt myoblasts with doxycycline-induced P3F expression. Transduction of the miR-17-92 construct resulted in high-level expression of pri-miR-17-92 and mature miR-17-92 miRNAs (Figure S4B, S4C) and modestly enhanced proliferation (Figure 5D) and clonogenic colony formation (Figure 5E) but did not induce focus formation (Figure 5F). These findings indicate that, while miR-17-92 is necessary for transformation when inhibited and can augment certain growth-related features when added, this miRNA cluster can neither substitute for a primary transformation signal nor can it function similar to MYCN in collaborating with P3F to transform cells. These findings underscore the requirement for additional oncogenic signals or cooperating genetic events to achieve full oncogenic transformation.

### A Distal P3F DNA-Binding Element Regulates *MIR17HG* Expression

To investigate the transcriptional regulation of *MIR17HG* by P3F in FP-RMS, we analyzed P3F chromatin immunoprecipitation sequencing (ChIP-seq) datasets from Rh4 FP-RMS cells (26) and Dbt-iP3F myoblasts (27). These analyses revealed a prominent P3F-bound DNA element in Rh4 cells located approximately 1.84 Mb upstream of the MIR17HG locus (Figure 6A, S5A). This same site was also detected 24 hours following Dox-induced P3F expression in Dbt-iP3F myoblasts. Consistent with previous findings that P3F often regulates transcription through distal intergenic enhancers, published datasets indicate that this region also displayed a MED1 co-activator binding peak and high H3K27ac enrichment, which are hallmarks of active enhancer complexes (26–29). Notably, there were no other recurrent P3F- or MED1-bound sites between this enhancer and the *MIR17HG* gene in both cell lines.

**Figure 6.**
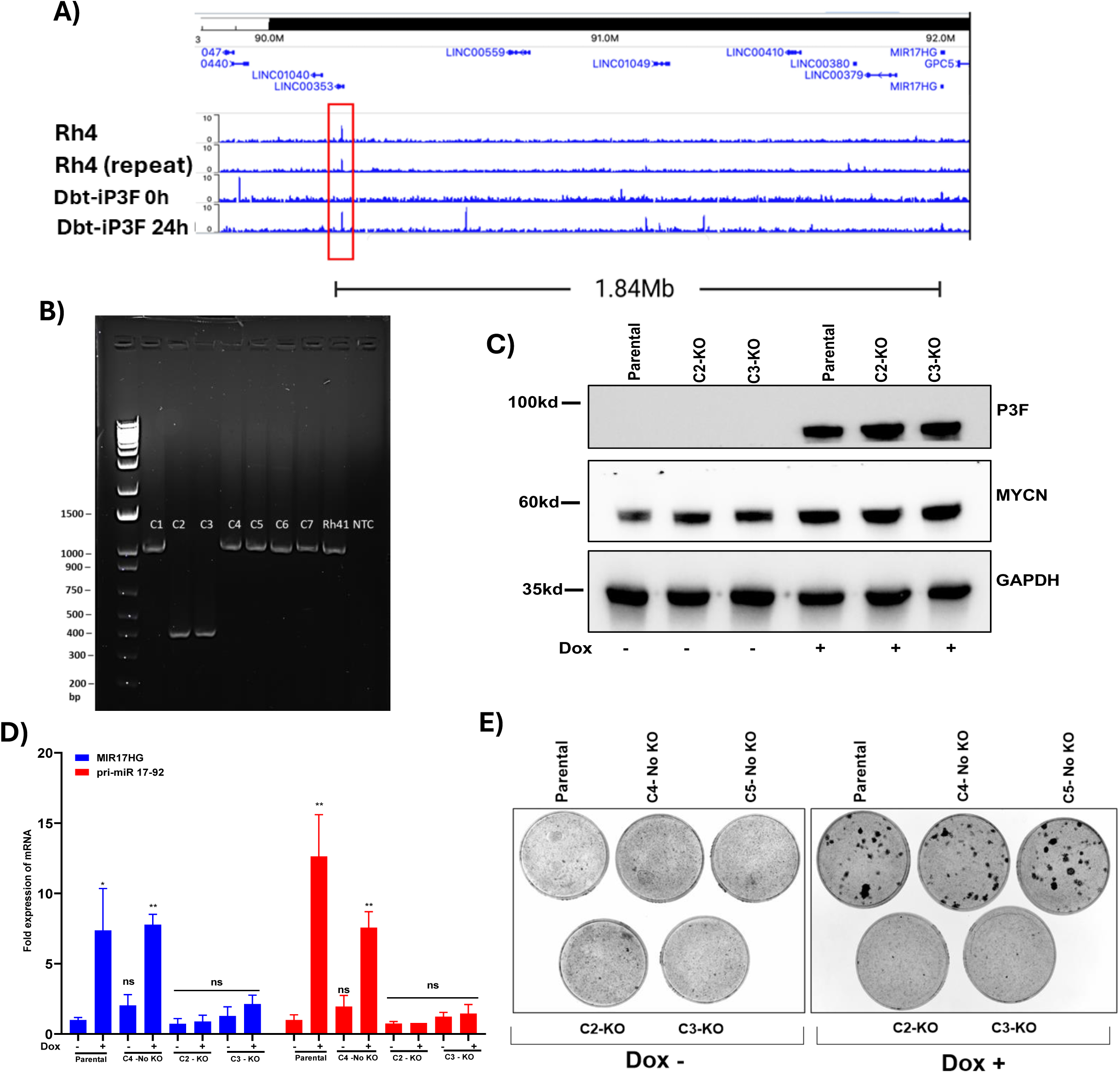
A distal P3F-binding element regulates *MIR17HG* expression and oncogenic transformation. **A.** Genomic map showing P3F ChIP-seq enrichment in Rh4 and in Dbt-iP3F cells at 0 and 24-hours following treatment with 500 ng/ml doxycycline (Dox). Data were obtained from published datasets [20, 21]. **B.** Agarose gel assaying Dbt-MYCN-iP3F subclones (C1-C7) for deletion of upstream region following treatment with gRNAs flanking distal P3F-binding site. **C.** Western blot analysis of P3F and MYCN protein in parental cells and deleted (C2, C3) subclones. GAPDH serves as a loading control. **D.** qPCR analysis of *MIR17HG* and pri-miR-17-92 transcript levels in deleted (C2, C3) subclones compared to parental and non-deleted (C4) controls. Results are normalized for parental cells without Dox. Data are presented as mean ± SD from at least three independent experiments. Statistical significance is displayed as described in Figure 1. **(E)** Focus formation assay of deleted (C2, C3) subclones compared to parental and non-deleted (C4, C5) controls. In parts C, D and E, cells were treated without (-) or with (+) 500 ng/ml Dox.

To assess the functional significance of this distal binding site, we utilized a CRISPR/Cas9-mediated strategy to delete this site in Dbt-MYCN-iP3F myoblasts. This deletion was identified in two subclones (C2 and C3) (Figure 6B) and was validated by Sanger sequencing of PCR products (Figure S5B). Western blot analysis confirmed that P3F and MYCN protein levels remained unchanged following deletion (Figure 6C), indicating that the observed effects were not due to loss of these oncogenic transcription factors. Expression analysis by qPCR demonstrated a significant reduction in *MIR17HG*, pri-miR-17-92 and mature miR-17-92 miRNA levels in the deleted subclones compared to parental and undeleted subclones (Figure 6D, S5C). These findings suggest that this distal P3F-bound region is critical for transcriptional activation of the *MIR17HG* locus.

At the phenotypic level, the deleted subclones showed a marked reduction in focus-forming ability indicative of a loss of oncogenic transformation (Figure 6E). Despite the loss of transformation, live-cell imaging revealed that these deleted subclones do not lose proliferative activity (data not shown), suggesting that this regulatory element is dispensable for basal growth but necessary for transformation-associated phenotypes.

To determine whether this P3F-binding site regulates other nearby transcripts, we analyzed RNA-seq data (from parental Dbt-MYCN-iP3F cells with and without Dox) for the expression of surrounding genes, including long intergenic non-coding RNAs, miRNAs and protein-coding genes within a 2 Mb window around this P3F binding site (Figure S5A). Surprisingly, none of these genes showed detectable expression in the RNA-seq dataset, except for *MIR17HG*. The complete absence of expression of neighboring loci supports the specificity of this regulatory element in controlling *MIR17HG* transcription. These findings provide compelling evidence that the identified distal P3F-binding site serves as a regulatory element critical for *MIR17HG* expression in FP-RMS.

## Discussion

The *MIR17HG* locus contains the polycistronic miR-17-92 cluster, which comprises six miRNAs: miR-17, miR-18a, miR-19a, miR-20a, miR-19b-1 and miR-92a-1 (30). This cluster is a well-established oncomir, which is amplified (31–33) or overexpressed (22) across a wide range of human cancers, and contributes to malignant progression by enhancing proliferation, inhibiting apoptosis, and promoting angiogenesis and metastasis (34). In colon cancer, members of the miR-17-92 cluster activate Wnt/β-catenin signaling and epithelial-mesenchymal transition (35), while suppressing TGF-β signaling via direct targeting of *TGFBR2*, *VEGFA* and *HIF1A* (36). In renal cell carcinoma, overexpression of miR-17 and miR-20a enhances cell proliferation by modulating the HIF and mTOR pathways (37). In breast cancer, the cluster promotes proliferation and invasion, particularly in triple-negative subtypes through repression of HBP1 (38) and activation of Wnt signaling (39). In pediatric malignancies such as neuroblastoma and medulloblastoma, *MYCN* amplification is a hallmark genetic feature and has been shown to drive *MIR17HG* expression (40,41). In these pediatric tumors, MYCN activates transcription of the miR-17-92 cluster, contributing to tumorigenesis by suppressing differentiation and apoptosis, reinforcing its role as a downstream effector of MYCN-driven tumor biology (41).

In this study, we focused on the regulation and function of the miR-17-92 cluster in the pediatric soft tissue cancer FP-RMS. Our previous observations of FP-RMS tumors noted that 13q31 amplification occurs in a subset of FP-RMS, most notably in tumors harboring the *P7F* fusion. Array-based copy number studies localized *MIR17HG* to the minimal common region of 13q31 amplification. Although there is generally increased expression of the miR-17-92 cluster in cases with amplification, miR-17-92 expression does not strictly correlate with amplification status. A substantial fraction of FP-RMS tumors lacking 13q31 amplification, particularly among *P3F*-positive cases, show high expression, pointing to the presence of amplification-independent regulatory mechanisms (10). We address this issue by defining a cooperative transcriptional mechanism through which the oncogenic fusion protein P3F and the MYCN transcription factor drive expression of *MIR17HG* in FP-RMS. This cooperative interaction is essential for establishing and maintaining the transformed phenotype of FP-RMS cells, implicating *MIR17HG* as a critical downstream oncogenic effector of the P3F-MYCN axis.

Chromatin immunoprecipitation studies identified a regulatory element located 1.84 Mb upstream of the *MIR17HG* locus (26). Our studies indicate that this element is required for transcriptional activation of the *MIR17HG* gene. While there is no evidence of DNA looping of this region to the *MIR17HG* locus, this region harbors a P3F binding motif and is enriched for P3F binding by ChIP-seq. Furthermore, deletion of this region abrogates *MIR17HG* activation, even in the presence of P3F and MYCN. We therefore propose that this element functions as a distal enhancer element contributing to *MIR17HG* transcription in FP-RMS and provides the mechanistic basis for high *MIR17HG* expression in non-amplified FP-RMS cases.

Our findings raise important questions about regulatory differences between the *P3F*- and *P7F*-positive subtypes of FP-RMS. Our previous studies demonstrated that both *P3F* and *P7F* fusion genes are overexpressed relative to their respective wild-type counterparts, *PAX3* and *PAX7*; however, the mechanisms underlying their overexpression are distinct. Specifically, *P3F* is typically upregulated through a copy number-independent transcriptional mechanism, while *P7F* is more often overexpressed due to genomic amplification (42). We now propose that there are two different mechanisms to overexpress the *MIR17HG* locus, a copy number-independent transcriptional process that occurs predominantly in *P3F*-positive tumors and a genomic amplification process that occurs predominantly in *P7F*-positive tumors (10). These findings thus suggest that, for both the fusion gene and the *MIR17HG* gene, amplification may serve as an alternative mechanism in *P7F*-positive tumors to upregulate expression. This difference between the *P3F*- and *P7F*-positive subtypes cannot yet be fully explored because of the absence of suitable *P7F*-positive RMS cell line models with *P7F* and/or *MIR17HG* amplification. Nevertheless, our findings establish a framework for exploring how genomic and transcriptional mechanisms converge as alternative means to upregulate oncogene expression.

Collectively, our functional experiments demonstrate that the miR-17-92 cluster is required to sustain the oncogenic phenotype in FP-RMS. In particular, this miRNA cluster contributes to the proliferative and clonogenic activity of these FP-RMS cells, and without this proliferative stimulus, the oncogenic transformation phenotype is lost. In studies of two FP RMS lines and a myoblast model of FP-RMS, there is cell line-specific variability in the miRNAs that are necessary for proliferation and transformation. Regardless of the specific roles played by each miRNA in different cell lines, the combination of the six miRNAs in this cluster (or possibly even the combination of three miRNAs in SET1) provides a stimulatory signal, and this stimulatory combination thus points to the possibility of novel potential therapeutic targets in FP-RMS.

In summary, this study defines a previously unrecognized transcriptional circuit in FP-RMS by which P3F and MYCN cooperatively activate *MIR17HG* expression, promoting a miRNA program that enforces the oncogenic phenotype. These findings not only expand our understanding of how fusion oncoproteins regulate non-coding RNAs but also highlight the miR-17-92 cluster as a key effector and potential therapeutic vulnerability in FP-RMS. Therapeutic strategies that target this axis by disrupting enhancer activity, transcription factor binding or miRNA function can offer new avenues for treating this aggressive pediatric cancer.

## Acknowledgments

This research was supported by the Intramural Research Program of the National Institutes of Health (NIH) and the Joanna McAfee Childhood Cancer Foundation. The contributions of the NIH authors were made as part of their official duties as NIH federal employees, are in compliance with agency policy requirements, and are considered Works of the United States Government. However, the findings and conclusions presented in this paper are those of the authors and do not necessarily reflect the views of the NIH or the U.S. Department of Health and Human Services.

## Supplementary Figures

**Figure S1.**
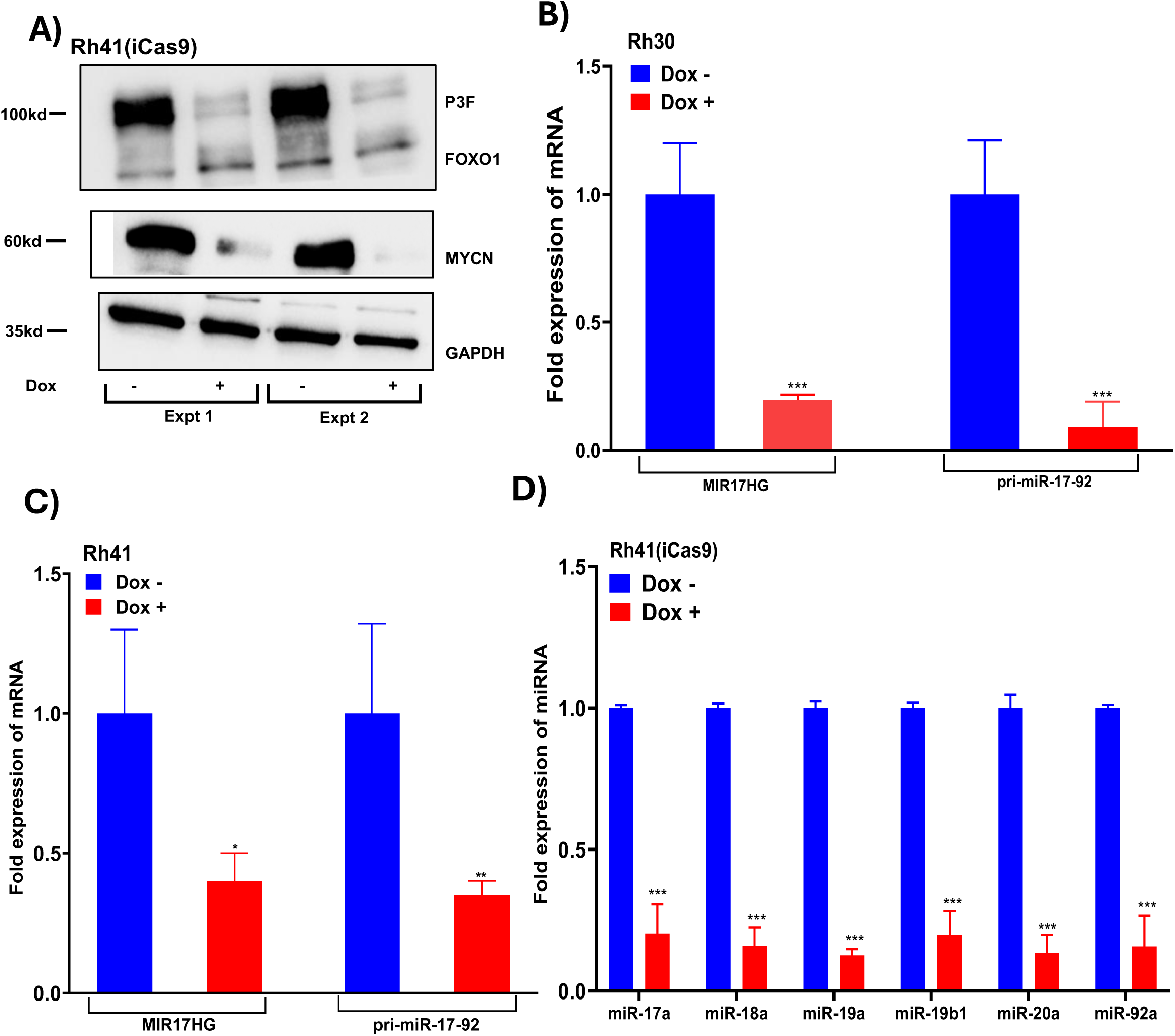
CRISPR-Cas9 knockout of P3F suppresses *MIR17HG* transcript, pri-miR-17-92 and mature miRNAs in FP-RMS cells. **A.** Western blot of P3F and MYCN protein in Rh41 cells treated without (-) or with (+) 2000 ng/ml doxycycline (Dox) to express Cas9. Two independent experiments are shown. **B, C.** qPCR analysis of *MIR17HG* transcript and pri-miR-17-92 in Rh30 (B) and Rh41 (C) following Dox-induced knockout of *P3F* in the Cas9-inducible subclones. **D**. qPCR analysis of mature miRNAs within the miR-17-92 cluster following Dox-induced knockout of *P3F* in the Cas9-inducible Rh41 cells. In B, C and D, cells were treated without (-) and with (+) Dox, and results are normalized for cells without Dox (-). Statistical significance is displayed as: not significant (ns), p < 0.05 (*), p < 0.01 (**), p < 0.001 (***).

**Figure S2.**
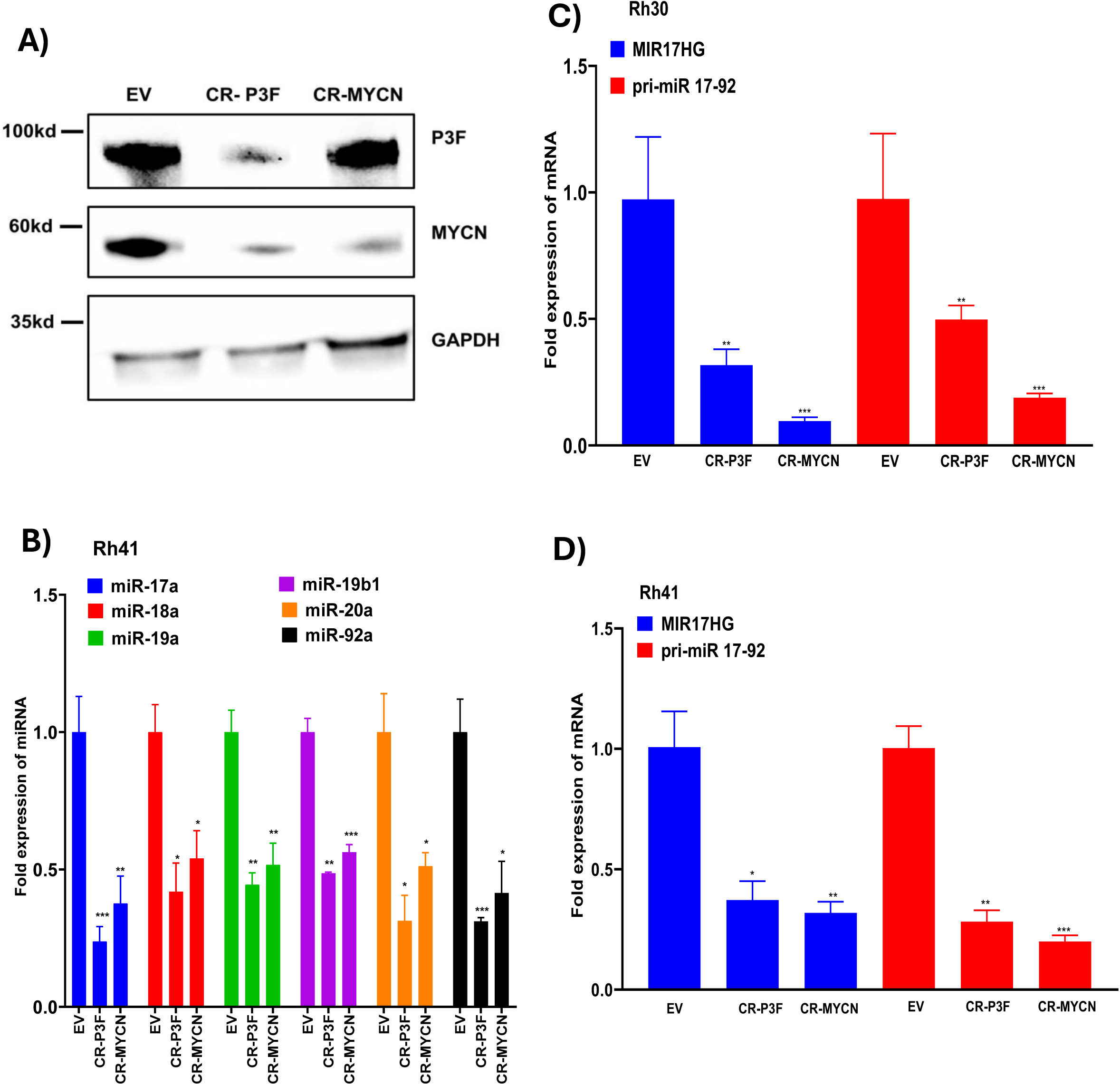
CRISPR-Cas9 knockout of P3F or MYCN suppresses MIR17HG transcript, pri-miR-17-92 and miR-17-92 cluster expression in FP-RMS cells. **A, B.** Western blot analysis of P3F and MYCN (A) and qPCR analysis of expression of mature miRNAs from the miR-17-92 cluster (B) following CRISPR-Cas9-mediated knockout (CR) of *P3F* or *MYCN* in Rh41 cells. Control cells were transduced with empty vector (EV). **C, D.** qPCR analysis of expression of *MIR17HG* transcript and pri-miR-17-92 following CRISPR-Cas9-mediated knockout of *P3F* or *MYCN* in Rh30 (C) and Rh41 (D) cells. In B, C, and D, results are normalized for control cells and are presented as mean ± SD from at least three independent experiments. Statistical significance is displayed as described in Figure S1.

**Figure S3.**
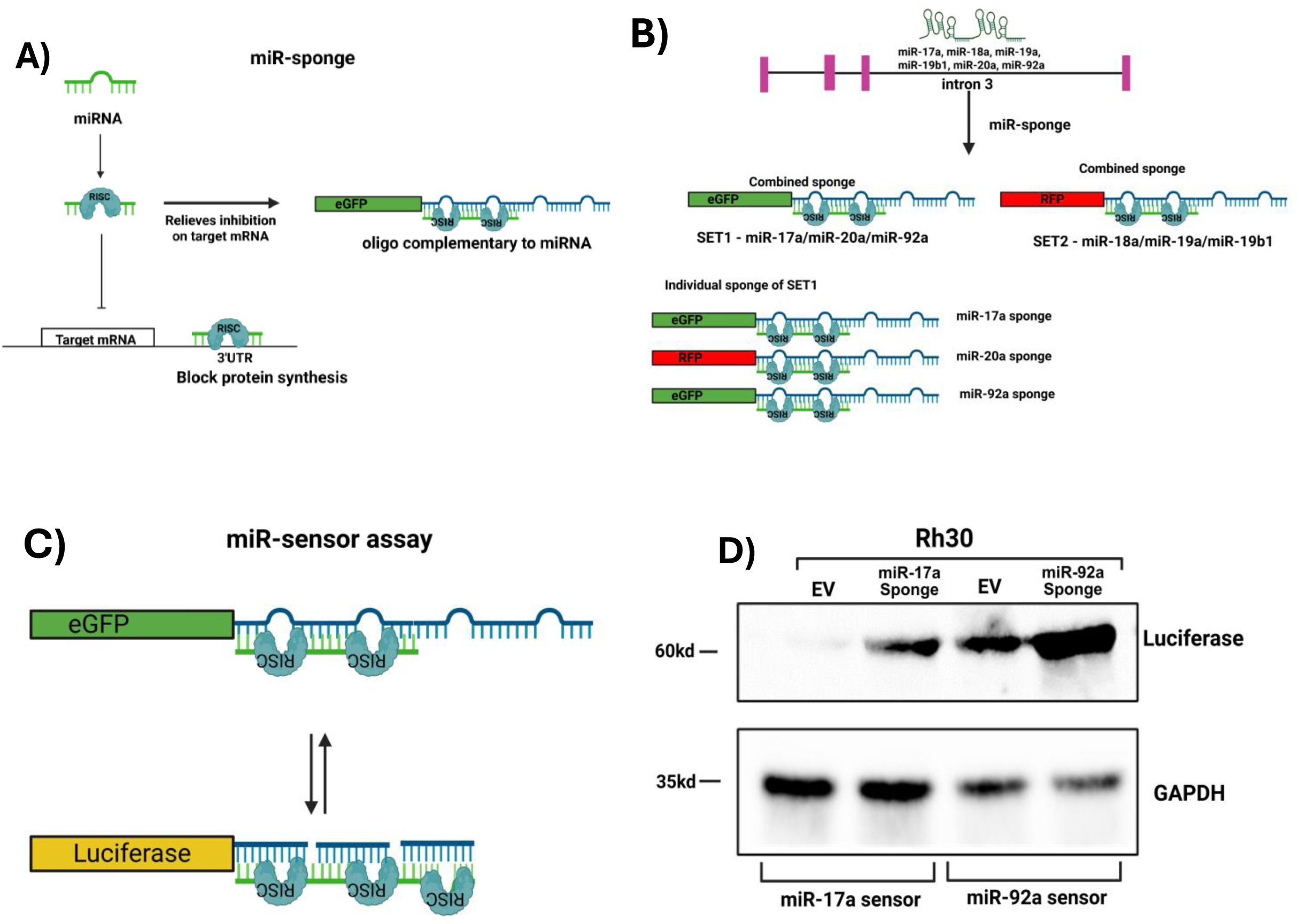
Inhibition of miR-17-92 with miRNA-sponges. **A**. Schematic representation of miRNA-sponge mechanism, illustrating synthetic constructs with tandem binding sites partially complementary to miRNA designed to sequester endogenous miRNAs and relieve repression of target mRNAs. **B**. Schematic of miRNA sponge design for the miR-17-92 cluster. The six miRNAs encoded within *MIR17HG* intron 3 were divided into two sponge constructs: SET1 (eGFP; miR-17a/miR-20a/miR-92a) and SET2 (RFP; miR-18a/miR-19a/miR-19b1). SET1 was further separated into individual sponges targeting miR-17a (eGFP), miR-20a (RFP), and miR-92a (eGFP). **C**. Schematic of the miRNA sensor assay. Sensor vectors contain three tandem binding sites with perfect complementarity to miR-17a or miR-92a placed in the 3′ UTR of a luciferase reporter gene. **D**. Western blot analysis of luciferase protein in doxycycline-treated Rh30 cells transduced with miR-17a sensor with EV or miR-17a sponge, or miR-92a sensor with EV or miR-92a sponge. GAPDH was utilized as a loading control.

**Figure S4.**
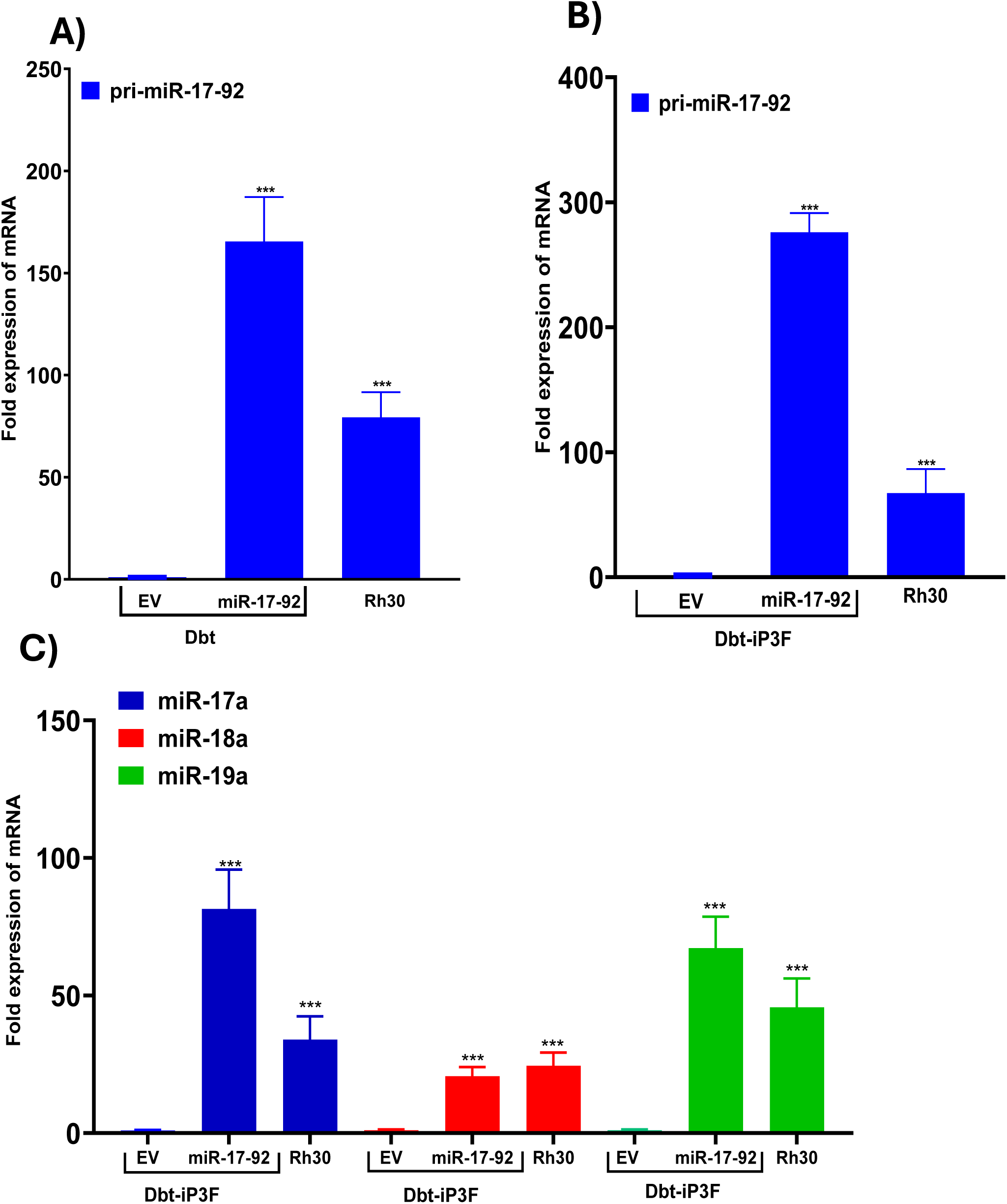
Effects of miR-17-92 expression in Dbt wild-type and Dbt-iP3F cells. **A.** qPCR analysis of pri-miR-17-92 expression in wild-type Dbt cells transduced with miR-17-92-containing construct or empty vector (EV) control. Data are normalized to 18S RNA and presented as mean ± SD from three replicates. **B, C.** qPCR analysis of pri-miR-17-92 (B) and mature miRNAs (C) encoded by the miR-17-92 cluster in Dbt-iP3F cells transduced with miR-17-92-containing construct or EV. Values are normalized to 18S and RNU6, respectively and presented as mean ± SD. Statistical significance is displayed as described in Figure S1.

**Figure S5.**
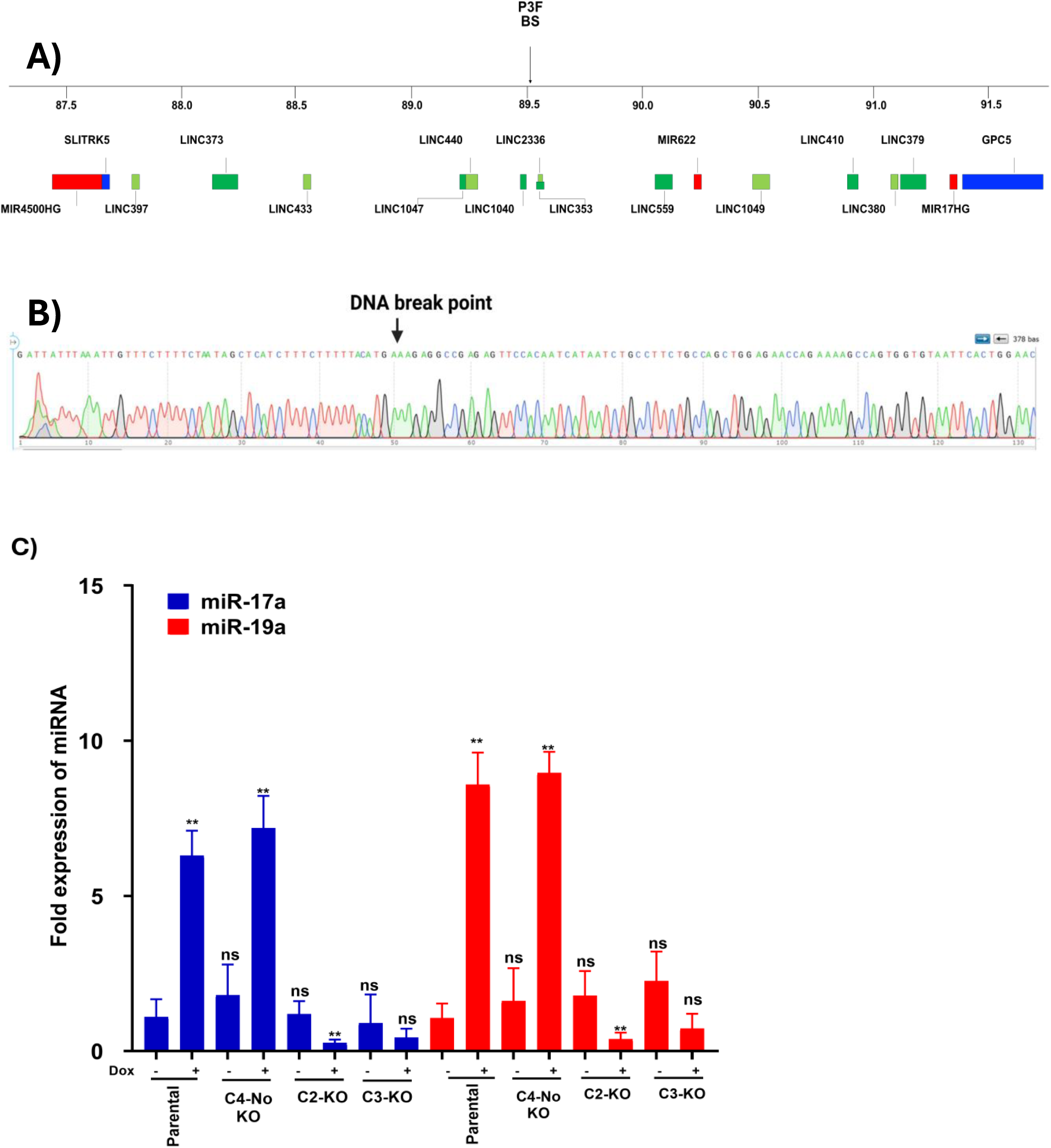
Identification of a distal P3F-binding site upstream of *MIR17HG*. **A.** Genomic map of region upstream of *MIR17HG* gene. The P3F binding site (BS) is shown above the horizontal line and distances (in Mb) are shown below the line. The horizontal boxes represent genes in this region (protein-coding, blue; miRNA-containing, red; long noncoding RNA, green). **B.** Sequencing chromatogram of deletion breakpoint in deleted subclones C2 and C3. **C.** qPCR analysis of miR-17a and miR-19a expression in deleted (C2, C3) subclones compared to parental and non-deleted (C4) controls. Cells were treated without (-) or with (+) 500 ng/ml doxycycline (Dox). Results are normalized for parental cells without Dox. Statistical significance is displayed as described in Figure S1.

## References

1. Barr FG, Galili N, Holick J, Biegel JA, Rovera G, Emanuel BS. Rearrangement of the PAX3 paired box gene in the paediatric solid tumour alveolar rhabdomyosarcoma. Nat Genet 1993;3:113–7

2. Davis RJ, D’Cruz CM, Lovell MA, Biegel JA, Barr FG. Fusion of PAX7 to FKHR by the variant t(1;13)(p36;q14) translocation in alveolar rhabdomyosarcoma. Cancer Res 1994;54:2869–72

3. Khan J, Bittner ML, Saal LH, Teichmann U, Azorsa DO, Gooden GC, et al. cDNA microarrays detect activation of a myogenic transcription program by the PAX3-FKHR fusion oncogene. Proc Natl Acad Sci USA 1999;96:13264–9

4. Cao L, Yu Y, Bilke S, Walker RL, Mayeenuddin LH, Azorsa DO, et al. Genome-wide identification of PAX3-FKHR binding sites in rhabdomyosarcoma reveals candidate target genes important for development and cancer. Cancer Res 2010;70:6497–508

5. Ha M, Kim VN. Regulation of microRNA biogenesis. Nat Rev Mol Cell Biol 2014;15:509–24

6. Hong L, Lai M, Chen M, Xie C, Liao R, Kang YJ, et al. The miR-17-92 cluster of microRNAs confers tumorigenicity by inhibiting oncogene-induced senescence. Cancer Res 2010;70:8547–57

7. Matsubara H, Takeuchi T, Nishikawa E, Yanagisawa K, Hayashita Y, Ebi H, et al. Apoptosis induction by antisense oligonucleotides against miR-17-5p and miR-20a in lung cancers overexpressing miR-17-92. Oncogene 2007;26:6099–105

8. Ernst A, Campos B, Meier J, Devens F, Liesenberg F, Wolter M, et al. De-repression of CTGF via the miR-17-92 cluster upon differentiation of human glioblastoma spheroid cultures. Oncogene 2010;29:3411–22

9. Farazi TA, Spitzer JI, Morozov P, Tuschl T. miRNAs in human cancer. J Pathol 2011;223:102–15

10. Reichek JL, Duan F, Smith LM, Gustafson DM, O’Connor RS, Zhang C, et al. Genomic and clinical analysis of amplification of the 13q31 chromosomal region in alveolar rhabdomyosarcoma: a report from the Children’s Oncology Group. Clin Cancer Res 2011;17:1463–73

11. Rota R, Ciarapica R, Giordano A, Miele L, Locatelli F. MicroRNAs in rhabdomyosarcoma: pathogenetic implications and translational potentiality. Mol Cancer 2011;10:120

12. Schulte JH, Horn S, Otto T, Samans B, Heukamp LC, Eilers UC, et al. MYCN regulates oncogenic MicroRNAs in neuroblastoma. Int J Cancer 2008;122:699–704

13. Marshall AD, Grosveld GC. Alveolar rhabdomyosarcoma - The molecular drivers of PAX3/7-FOXO1-induced tumorigenesis. Skelet Muscle 2012;2:25

14. Barta T, Peskova L, Hampl A. miRNAsong: a web-based tool for generation and testing of miRNA sponge constructs in silico. Sci Rep 2016;6:36625

15. Krüger J, Rehmsmeier M. RNAhybrid: microRNA target prediction easy, fast and flexible. Nucleic Acids Res 2006;34:W451–4

16. Zhang X, Zhou Y, Chen S, Li W, Chen W, Gu W. LncRNA MACC1-AS1 sponges multiple miRNAs and RNA-binding protein PTBP1. Oncogenesis 2019;8:73

17. Boudjadi S, Pandey PR, Chatterjee B, Nguyen TH, Sun W, Barr FG. A Fusion Transcription Factor-Driven Cancer Progresses to a Fusion-Independent Relapse via Constitutive Activation of a Downstream Transcriptional Target. Cancer Res 2021;81:2930–42

18. Nguyen TH, Vemu PL, Hoy GE, Boudjadi S, Chatterjee B, Shern JF, et al. Serine hydroxymethyltransferase 2 expression promotes tumorigenesis in rhabdomyosarcoma with 12q13-q14 amplification. J Clin Invest 2021;131: e138022

19. Schneider CA, Rasband WS, Eliceiri KW. NIH Image to ImageJ: 25 years of image analysis. Nat Methods 2012;9:671–5

20. Zargar S, Tomar V, Shyamsundar V, Vijayalakshmi R, Somasundaram K, Karunagaran D. A Feedback Loop between MicroRNA 155 (miR-155), Programmed Cell Death 4, and Activation Protein 1 Modulates the Expression of miR-155 and Tumorigenesis in Tongue Cancer. Mol Cell Biol 2019;39:e00410–18

21. Kumps C, Fieuw A, Mestdagh P, Menten B, Lefever S, Pattyn F, et al. Focal DNA copy number changes in neuroblastoma target MYCN regulated genes. PLoS One 2013;8:e52321

22. Northcott PA, Fernandez LA, Hagan JP, Ellison DW, Grajkowska W, Gillespie Y, et al. The miR-17/92 polycistron is up-regulated in sonic hedgehog-driven medulloblastomas and induced by N-myc in sonic hedgehog-treated cerebellar neural precursors. Cancer Res 2009;69:3249–55

23. Pandey PR, Chatterjee B, Olanich ME, Khan J, Miettinen MM, Hewitt SM, et al. PAX3-FOXO1 is essential for tumour initiation and maintenance but not recurrence in a human myoblast model of rhabdomyosarcoma. J Pathol 2017;241:626–37

24. Jung J, Yeom C, Choi YS, Kim S, Lee E, Park MJ, et al. Simultaneous inhibition of multiple oncogenic miRNAs by a multi-potent microRNA sponge. Oncotarget 2015;6:20370–87

25. Ebert MS, Sharp PA. MicroRNA sponges: progress and possibilities. RNA 2010;16:2043–50

26. Gryder BE, Yohe ME, Chou HC, Zhang X, Marques J, Wachtel M, et al. PAX3-FOXO1 Establishes Myogenic Super Enhancers and Confers BET Bromodomain Vulnerability. Cancer Discov 2017;7:884–99

27. Sunkel BD, Wang M, LaHaye S, Kelly BJ, Fitch JR, Barr FG, et al. Evidence of pioneer factor activity of an oncogenic fusion transcription factor. iScience 2021;24:102867

28. Kucinski J, Tallan A, Taslim C, Vontell AM, Silvius KM, Wang M, et al. Rhabdomyosarcoma fusion oncoprotein initially pioneers a neural signature in vivo. Cell Reports 2025;44:115923

29. Cao L, Yu Y, Bilke S, Walker RL, Mayeenuddin LH, Azorsa DO, et al. Genome-wide identification of PAX3-FKHR binding sites in rhabdomyosarcoma reveals candidate target genes important for development and cancer. Cancer Res 2010;70:6497–508

30. He L, Thomson JM, Hemann MT, Hernando-Monge E, Mu D, Goodson S, et al. A microRNA polycistron as a potential human oncogene. Nature 2005;435:828–33

31. Martens-de Kemp SR, Komor MA, Hegi R, Bolijn AS, Tijssen M, de Groen FLM, et al. Overexpression of the miR-17-92 cluster in colorectal adenoma organoids causes a carcinoma-like gene expression signature. Neoplasia 2022;32:100820

32. Diosdado B, van de Wiel MA, Terhaar Sive Droste JS, Mongera S, Postma C, Meijerink WJHJ, et al. MiR-17-92 cluster is associated with 13q gain and c-myc expression during colorectal adenoma to adenocarcinoma progression. Brit J Cancer 2009;101:707–14

33. Ota A, Tagawa H, Karnan S, Tsuzuki S, Karpas A, Kira S, et al. Identification and characterization of a novel gene, C13orf25, as a target for 13q31-q32 amplification in malignant lymphoma. Cancer Res 2004;64:3087–95

34. Olive V, Jiang I, He L. mir-17-92, a cluster of miRNAs in the midst of the cancer network. Int J Biochem Cell Biol 2010;42:1348–54

35. Werner H. Tumor suppressors govern insulin-like growth factor signaling pathways: implications in metabolism and cancer. Oncogene 2012;31:2703–14

36. Taguchi A, Yanagisawa K, Tanaka M, Cao K, Matsuyama Y, Goto H, et al. Identification of hypoxia-inducible factor-1 alpha as a novel target for miR-17-92 microRNA cluster. Cancer Res 2008;68:5540–5

37. Chow TF, Mankaruos M, Scorilas A, Youssef Y, Girgis A, Mossad S, et al. The miR-17-92 cluster is over expressed in and has an oncogenic effect on renal cell carcinoma. J Urol 2010;183:743–51

38. Li H, Bian C, Liao L, Li J, Zhao RC. miR-17-5p promotes human breast cancer cell migration and invasion through suppression of HBP1. Breast Cancer Res Treat 2011;126:565–75

39. Zhao W, Gupta A, Krawczyk J, Gupta S. The miR-17-92 cluster: Yin and Yang in human cancers. Cancer Treat Res Commun 2022;33:100647

40. O’Donnell KA, Wentzel EA, Zeller KI, Dang CV, Mendell JT. c-Myc-regulated microRNAs modulate E2F1 expression. Nature 2005;435:839–43

41. Schulte JH, Horn S, Otto T, Samans B, Heukamp LC, Eilers UC, et al. MYCN regulates oncogenic MicroRNAs in neuroblastoma. International journal of cancer 2008;122:699–704

42. Shern JF, Chen L, Chmielecki J, Wei JS, Patidar R, Rosenberg M, et al. Comprehensive genomic analysis of rhabdomyosarcoma reveals a landscape of alterations affecting a common genetic axis in fusion-positive and fusion-negative tumors. Cancer Discov 2014;4:216–31

